# Impaired functional connectivity of the hippocampus in murine models of NMDA-receptor antibody associated pathology

**DOI:** 10.1101/2022.01.12.476037

**Authors:** Joseph Kuchling, Betty Jurek, Mariya Kents, Jakob Kreye, Christian Geis, Jonathan Wickel, Susanne Mueller, Stefan Paul Koch, Philipp Boehm-Sturm, Harald Prüss, Carsten Finke

**Affiliations:** Department of Neurology and Experimental Neurology, Charité – Universitätsmedizin Berlin, corporate member of Freie Universität Berlin and Humboldt-Universität zu Berlin, Germany; Neurocure Cluster of Excellence, NeuroCure Clinical Research Center, Charité – Universitätsmedizin Berlin, corporate member of Freie Universität Berlin and Humboldt-Universität zu Berlin, Germany; Berlin Institute of Health at Charité – Universitätsmedizin Berlin, Charitéplatz 1, 10117 Berlin, Germany; German Center for Neurodegenerative Diseases (DZNE) Berlin, Berlin, Germany; Section of Translational Neuroimmunology, Hans Berger Department of Neurology, Jena University Hospital, Jena, Germany; Neurocure Cluster of Excellence, Core Facility 7T Experimental MRIs, Charité – Universitätsmedizin Berlin, corporate member of Freie Universität Berlin and Humboldt-Universität zu Berlin, Germany; Berlin Center for Stroke Research, Charité – Universitätsmedizin Berlin, corporate member of Freie Universität Berlin and Humboldt-Universität zu Berlin, Germany; Humboldt-Universität zu Berlin, Berlin School of Mind and Brain, Berlin, Germany

**Author notes:** Equally contributing authors (Harald Prüss, Carsten Finke). Correspondence:* Carsten Finke, Department of Neurology, Charité - Universitätsmedizin Berlin, Charitéplatz 1, 10117 Berlin, Germany, or Harald Prüss, Department of Neurology, Charité - Universitätsmedizin Berlin, Charitéplatz 1, 10117 Berlin, Germany.

## Abstract

**Introduction:** While decreased hippocampal connectivity and disruption of functional networks are established MRI features in human anti-N-methyl-D-aspartate receptor (NMDAR) encephalitis, the underlying pathophysiology for brain network alterations remains poorly understood. Application of patient-derived monoclonal antibodies against the NR1 subunit of the NMDAR allows for the investigation of potential functional connectivity alterations in experimental murine NMDAR antibody disease models.

**Objective:** To explore functional connectivity changes in NR1 antibody mouse models using resting-state functional MRI (rs-fMRI).

**Methods:** Adult C57BL/6J mice (n=10) were intrathecally injected with a recombinant human NR1 antibody over 14 days and then studied using rs-fMRI at 7 Tesla. In addition, a newly established mouse model with *in utero* exposure to a human recombinant NR1 antibody characterized by a neurodevelopmental disorder (NR1-offspring) was investigated with rs-fMRI at the age of 8 weeks (n=15) and 10 months (n=14). Mice exposed to isotype-matched control antibodies served as controls. Independent component analysis (ICA) and dual regression analysis were performed to compare functional connectivity between NMDAR antibody mouse models and control mice.

**Results:** Adult NR1-antibody injected mice showed significantly impaired functional connectivity within the dentate gyrus of the left hippocampus in comparison to controls, resembling impaired hippocampal functional connectivity patterns observed in human patients with NMDAR encephalitis. Similarly, analyses showed significantly reduced functional connectivity in the dentate gyrus in NR1-offspring compared after 8 weeks, and impaired connectivity in the dentate gyrus and CA3 hippocampal subregion in NR1-offspring at the age of 10 months.

**Conclusion:** Functional connectivity changes within the hippocampus resulting from both direct application and *in utero* exposure to NMDAR antibodies can be modeled in experimental murine systems. With this translational approach, we successfully reproduced functional MRI alterations previously observed in human NMDAR encephalitis patients. Future experimental studies will identify the detailed mechanisms that cause functional network alterations and may eventually allow for non-invasive monitoring of disease activity and therapeutic effects in autoimmune encephalitis.

## Introduction

Anti-N-methyl-D-aspartate receptor (NMDAR) encephalitis is an autoimmune disorder caused by autoantibodies targeting the NR1 subunit of the NMDAR. Patients present with neuropsychiatric symptoms including psychosis, behavioural changes and memory deficits, as well as seizures, movement disorders and decreased levels of consciousness ^1^. Despite the often severe clinical course, routine brain MRI yields no abnormalities in most patients and therefore provides only limited diagnostic and prognostic value ^2,3^. By contrast, resting-state functional MRI (rs-fMRI) studies revealed disruption of functional connectivity of the hippocampus with the anterior default-mode network (DMN) that correlated with memory performance ^4^. Recent large-scale network analyses confirmed these findings and showed decoupling of the medial temporal lobe network from the DMN that correlated with the severity of memory deficits as well as disruption of fronto-parietal networks related to psychiatric symptoms ^5^. These observations suggest that NMDAR antibody-associated clinical and cognitive symptoms are linked to disturbance of functional brain networks ^4,5^. Predominant antibody-mediated dysfunction of the hippocampus is further substantiated by localized atrophy of hippocampal subfields (Finke et al., 2016), in line with the fact that the hippocampal formation contains the highest density of NMDARs in the brain ^6^. However, the exact pathophysiological relationship between NMDAR antibody exposure and impaired functional connectivity remains elusive and can only partially be addressed by studies in humans.

Recently, several mouse models of NMDAR encephalitis have been developed ^7–11^. We have shown that monoclonal human cerebrospinal fluid-derived antibodies against the NR1 subunit of the NMDAR derived from NMDAR encephalitis patients have direct pathogenicity by causing neuronal surface receptor downregulation and subsequent disruption of synaptic NMDAR currents ^12,13^. Furthermore, injection of monoclonal NR1 antibodies in pregnant maternal mice with subsequent fetal *in utero* exposure resulted in a neurodevelopmental disorder in the offspring with long-lasting neuropathological effects reflected by increased postnatal mortality, hyperactivity, lower anxiety and impaired sensorimotor gating as well as reduced brain volumes ^14^. In a related model using patient IgG, brain changes such as thinning of cortical layers and behavior abnormalities were reversible until adulthood ^15^.

Recent advances in the acquisition and analysis of functional MRI data in mice now allow for robust and reproducible analysis of functional connectivity alterations and large-scale network dysfunction in mouse models of neurological disorders ^16,17^. This provides the unique opportunity to not only study the effect of NMDAR antibody exposure on the functional connectivity, but also to compare the connectivity changes from these animal models with the functional network alterations observed in human patients. Given that potential novel interventional strategies are preclinically first evaluated in mice before being tested in humans, studying the correlation between NMDAR antibody-mediated clinical and imaging features in mice may help to elucidate disease-specific markers that are related to the neuronal dysfunction.

Here, we performed 7T rs-fMRI investigations in mice that were injected with human recombinant NR1 antibodies and in the offspring of a recently established murine model with *in utero* exposure to NR1 antibodies. We hypothesized that (i) functional connectivity changes of the hippocampus known from human NMDAR encephalitis are similarly observable in NR1 antibody mouse models and that (ii) functional connectivity alterations are present in antibody-mediated neurodevelopmental brain disorders.

## Material and methods

### Animal experiments

Animal experiments were carried out in accordance with the ARRIVE guidelines ^18^, the EU Directive (2010/63/EU) for animal experiments and were approved by the local ethics committee for Animal Welfare (LaGeSO, Berlin, G0175/15; Thuringian state authorities, UKJ-17-053). Ten mice were directly exposed to NR1 antibodies during adulthood (NR1; cohort 1; Fig. 1). These C57BL/6J mice (age of 10 weeks) received bilateral infusion of 200 µg human monoclonal IgG1 antibody (high-affinity NR1-reactive [amino-terminal domain] IgG1 clone: #003-102 [n=10, “NR1”]) ^12,19^ in PBS for 14 days into the lateral ventricles using implanted catheters connected to osmotics pumps following established protocols ^20–22^. Ten mice received human monoclonal IgG1 antibody that is non-reactive to brain tissue (control clone: #mGO53) ^12,23^ into the lateral ventricles via implanted catheters and served as NR1-controls. MR investigation of NR1 mice and their respective controls was performed at the age of 12 weeks (see Figure 1).

**Figure 1.**
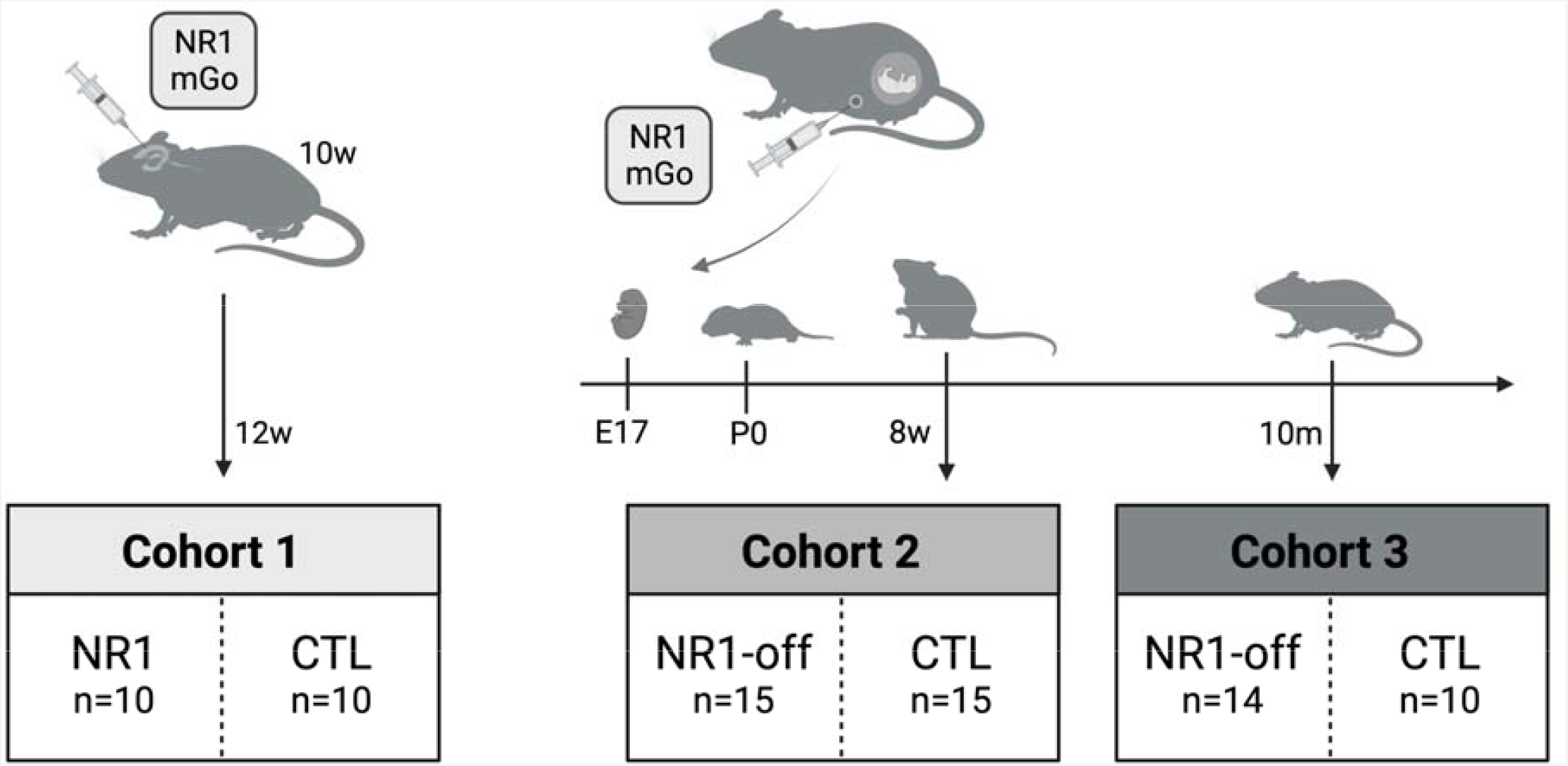
Flowchart of cohort selection. NR1 = NR1-reactive human monoclonal IgG1 antibody; mGo = monoclonal IgG1 antibody that is non-reactive to brain tissue (control clone: #mGO53); NR1-off = NR1-offspring cohorts; CTL = controls.

Additionally, 29 mice with prior *in utero* exposure to human recombinant NR1 antibody ^12,19^ were raised according to a previously published protocol (NR1-offspring; cohorts 2 and 3; Fig. 1) ^14^. In short, 8- to 10-week-old pregnant C57BL/6J mice were injected intraperitoneally, at gestational days E13 and E17, at each timepoint with 240 μg of human monoclonal IgG1 antibody #003-102. Twenty-five offspring mice from maternal animals exposed to intraperitoneally injected human monoclonal IgG1 antibody that is non-reactive to brain tissue (control clone: #mGO53) ^12,23^ (Fig. 1) served as controls for cohorts 2 and 3. All offspring mice were housed in treatment-mixed groups of 2-5 animals of both sexes and were investigated either at the age of 8 weeks (cohort 2; NR1-offspring: n=15; controls: n=15) or at the age of 10 months (cohort 3; NR1-offspring: n=14; controls: n=10) (see Figure 1).

### MRI data acquisition and quality control

Anesthesia was achieved using 1.5-2% isoflurane in a 70:30 nitrous oxide:oxygen mixture. Before start of the rsMRI scan, isoflurane levels were reduced to 1.2 (+/− 0.3) % until animals breathed regularly at higher rate of 160 (+/-25)/min. This process took 5-7 min and body temperature and respiration rate were monitored with MRI compatible equipment (Small Animal Instruments Inc., Stony Brook, NY). 2D echo-planar imaging (EPI) resting-state functional MR (rs-fMRI) images were acquired (repetition time (TR) = 1000 ms; echo time (TE) = 13 ms; flip angle (FA) = 50°; 300 repetitions; 16 axial slices with slice thickness = 0.75 mm; field of view (FOV) = 19.2 × 12.0 mm²; image matrix = 128 × 80; number of averages = 1) on a 7 T MR scanner (Bruker Biospec, Ettlingen, Germany) and a transmit/receive mouse cryoprobe (Bruker). Experimenters were blinded to the condition of the animals.

### rs-fMRI data processing and denoising

Processing of rs-fMRI data was modified according to an established rs-fMRI analysis protocol ^24^. Rs-fMRI datasets were skull-stripped using linear affine registration of an in-house mouse atlas template based on C57BL/6J control mice previously studied at the same scanner to individual mouse rs-fMRI images with FSL FLIRT ^25^. Each skull-stripped 4D dataset was then fed into FSL MELODIC ^26^ to perform within-subject spatial independent component analysis (ICA) with automatic dimensionality estimation, using a skull-stripped EPI image of a deliberately chosen subject as anatomical template. Within-subject ICA was performed after removal of the first 10 out of 300 time points of every subject and included in-plane smoothing with a 1.0 × 1.0 mm kernel. Subsequent denoising of single-subject fMRI independent components was performed in accordance with previously described protocols ^24,27^ by manual identification of signal and noise based on the threshold spatial maps, the temporal power spectrum, and the time course by one rater (JKu) who was blinded to the clinical phenotype of mice.

### Group ICA and dual regression

Resting state networks common to all mice were identified for each cohort, separately, by use of temporal-concatenation ICA with automatic component estimation as implemented in FSL MELODIC. To facilitate classification, resulting components were displayed as spatial color-coded z-maps onto the Allen Mouse Brain atlas (AMBA; mouse.brain-map.org/static/atlas) after co-registeration with FSL FLIRT. Group comparisons were carried out using dual regression and nonparametric permutation testing (1,000 permutations) with threshold-free cluster enhancement (TFCE) as implemented in FSL *randomise* (p < 0.05, familywise error [FWE]-corrected) ^28^. Resultant maps from group analyses were co-registered to AMBA for visualization.

## Results

### Resting State Component Identification

Group ICA identified canonical functional components with similar anatomical localization in cohort 1 (NR1 and controls; 90 components), cohort 2 (8w-offspring and 8w-controls; 90 components), and cohort 3 (10m-offspring and 10m-controls; 95 components). Component patterns consistently covered common neuroanatomical regions defined by co-registration with the AMBA, including brainstem, hypothalamus, thalamic areas with differentiation of thalamic subnuclei, somatosensory cortical areas, hippocampal formation and basal ganglia regions (Figure 2). Of note, the identification of functional components within the hippocampal formation allowed for the distinction between CA1-3, dentate gyrus and subiculum correlating with corresponding anatomical AMBA regions.

**Figure 2.**
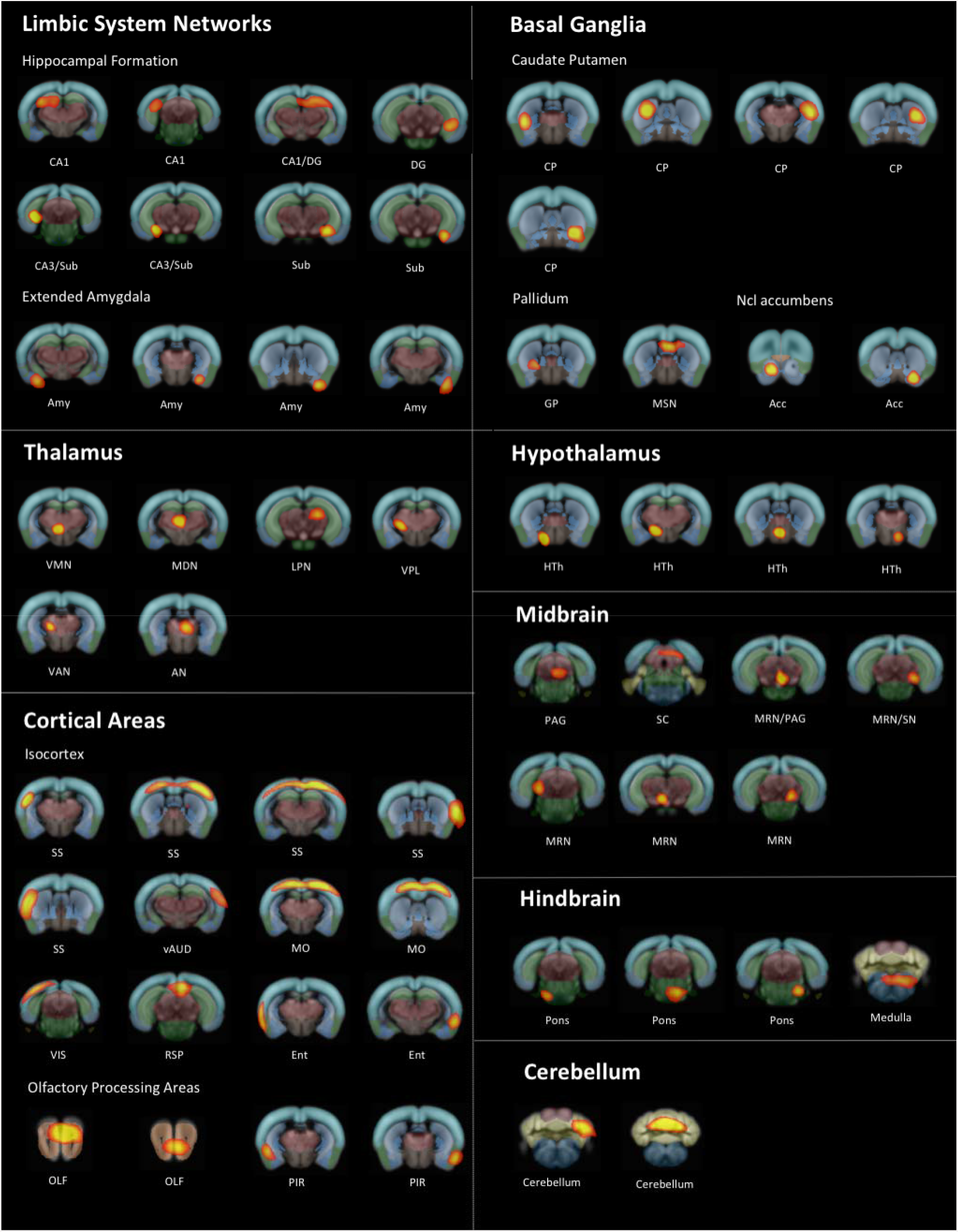
NR1 offspring at the age of 8 weeks and 8w-controls (cohort 2): Selection of brain components generated by group ICA. MELODIC group ICA analysis of 15 NR1 offspring and 15 healthy control mice at the age of 8 weeks generated anatomically plausible components displayed as spatial color-coded z-maps onto the AMBA. Components are arranged according to anatomical mouse brain structures after visual correlation with AMBA brain regions modified after previously published component grouping ^24,49^. DG = dentate gyrus; Sub = subiculum; Amy = amygdala; aCing = anterior cingulate area; VMN = thalamic ventromedial nucleus; LGN = thalamic lateral geniculate nucleus; MDN = medial dorsal nucleus; LPN = lateral posterior nucleus; VPL = thalamic ventral posterolateral nucleus; VAN = ventral anterior nucleus; AN = anterior nuclear group of the thalamus; SS = primary somatosensory area; vAUD = ventral auditory area; MO = pimary motor area; VIS = primary and anterolateral visual area; RSP = retrosplenial cortex; Ent = entorhinal area; Ent = lateral part of the entorhinal area; OLF = main olfactory bulb; PIR = piriform area; CP = caudate putamen; GP = globus pallidus; MSN = medial septal nucleus area of pallidum; Acc = nucleus accumbens; HTh = Hypothalamus; PAG = periaqueductal grey; SC = superior colliculus; MRN = midbrain

### Cohort 1: NR1 mice

NR1 mice showed significantly reduced functional connectivity in the left hippocampus in comparison to control mice (Figure 3). According to the AMBA, the major cluster of reduced functional connectivity (*p*=0.016) was primarily localized within left dentate gyrus. No further functional connectivity alterations were observed in NR1 mice compared to controls.

**Figure 3.**
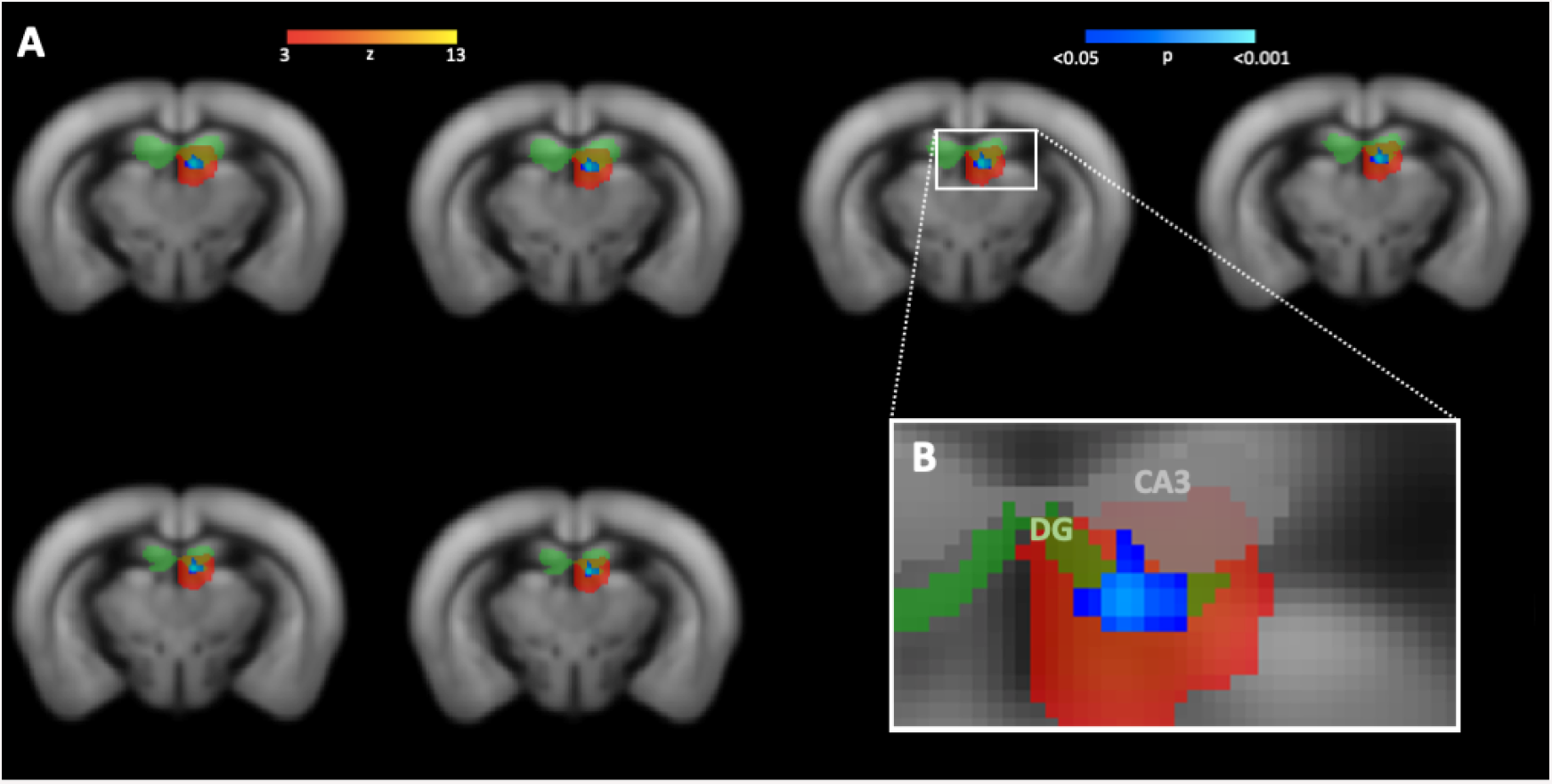
Reduced functional connectivity within hippocampus (dentate gyrus) of NR1 mice compared to controls (cohort 1). **A** Dual regression analysis revealed reduced functional connectivity of adult NR1 mice compared to controls (significantly different area displayed based on p-values in blue-lightblue) in a specific functional component (component area displayed as z-map in red-yellow). **B** Functional component and corresponding area of functional connectivity reduction is affiliated with left-hemispheric hippocampus (light-green region of interest) and **C** shows visual correspondence with hippocampal subfield of dentate gyrus (green). DG = dentate gyrus

### Cohort 2: 8w-offspring

Antibody-mediated effects of monoclonal NR1 antibodies in NR1 offspring mouse model investigated in cohorts 2 and 3 on the biomolecular and behavioural level have been previously described in detail at different developmental stages and have recently been published elsewhere ^14^. NR1-offspring at the age of 8 weeks exhibited selectively reduced functional connectivity of the left hippocampus (*p*=0.027) in comparison to control mice. Functional connectivity alterations were primarily located in the dentate gyrus according to the AMBA (Figure 4 A and B). No further significant functional connectivity differences between 8w-offspring mice and controls were observed.

**Figure 4.**
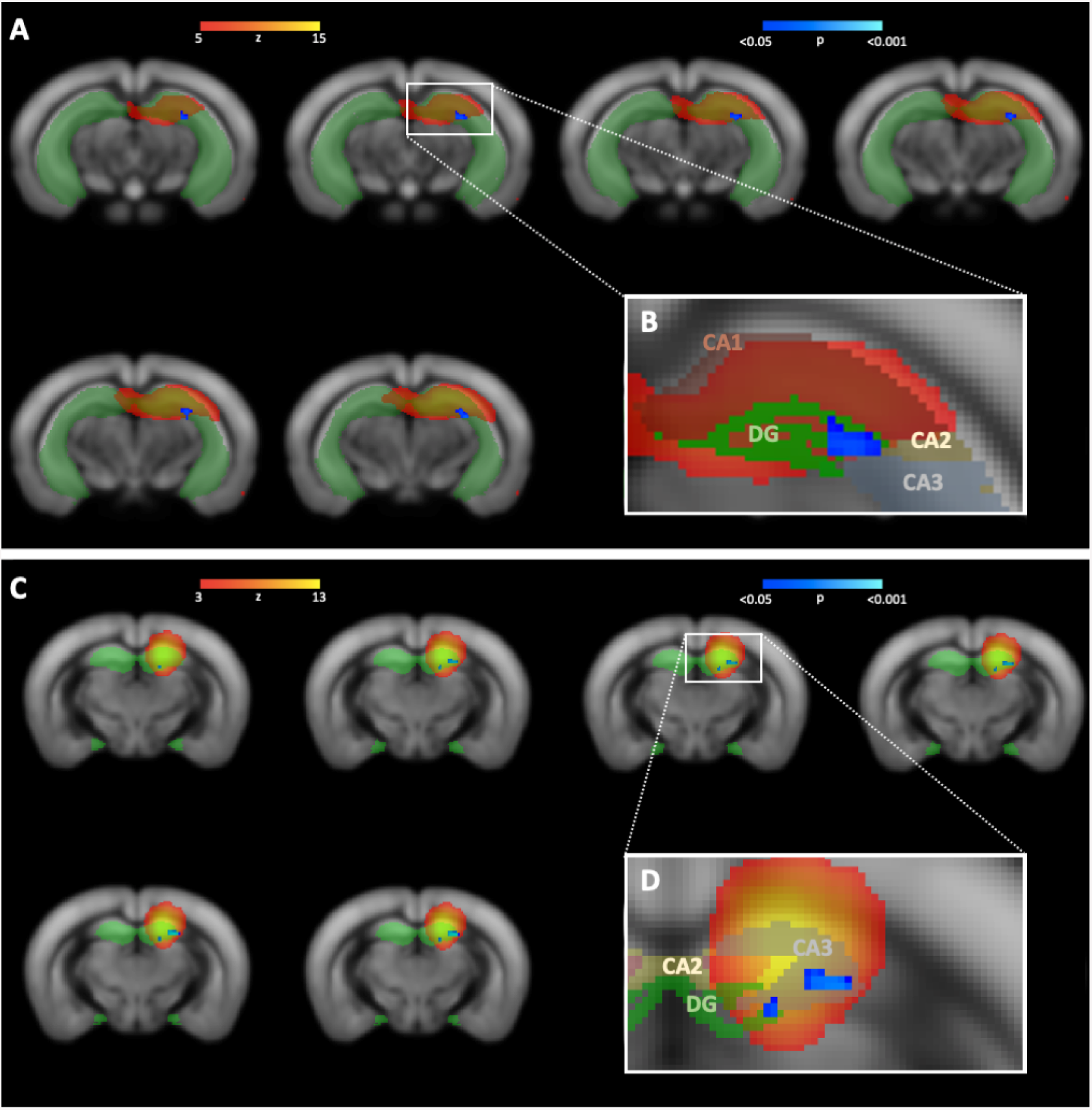
Reduced functional connectivity within hippocampus of NR1 offspring at different stages of age compared to controls (cohort 2 and 3). **A** Dual regression analysis revealed reduced functional connectivity of 8-week-old NR1 offspring with neurodevelopmental disorder compared to controls (significantly different area displayed based on p-values in blue-lightblue) in a specific functional component (component area displayed as z-map in red-yellow) affiliated with left-hemispheric hippocampus (light-green region of interest) and **B** shows visual correspondence with hippocampal subfield of dentate gyrus (green). **C** Reduced functional connectivity of 10-month-old NR1 offspring is detectable in a functional component corresponding with left-hemispheric hippocampus (light-green region of interest). **D** Visual correspondence of impaired connectivity cluster with hippocampal subfield of dentate gyrus (green) and CA3 (white). DG = dentate gyrus

### Cohort 3: 10m-offspring

Reduced functional connectivity was observed within the left hippocampus (*p*=0.010) in NR1-offspring at the age of 10 months compared to control mice. Specifically, functional connectivity alterations were located in the dentate gyrus and CA3 (Figure 4 C and D). No further functional connectivity differences between 10m-offspring mice and controls were found.

### Human NR1 antibodies bound to synaptic structures

Hippocampal staining revealed the well-known distribution pattern in pivotal regions of hippocampal connectivity, i.e. dentate gyrus and CA3 region, that receive projections from entorhinal cortex and from medial septum and contralateral hippocampus ^29^. Immunofluorescence of mouse brain sections post-MRI in cohort 1 confirmed the strong binding of intrathecally administered human monoclonal NR1 IgG antibodies on hippocampal neuropil on murine brain sections (Figure 5A).

**Figure 5.**
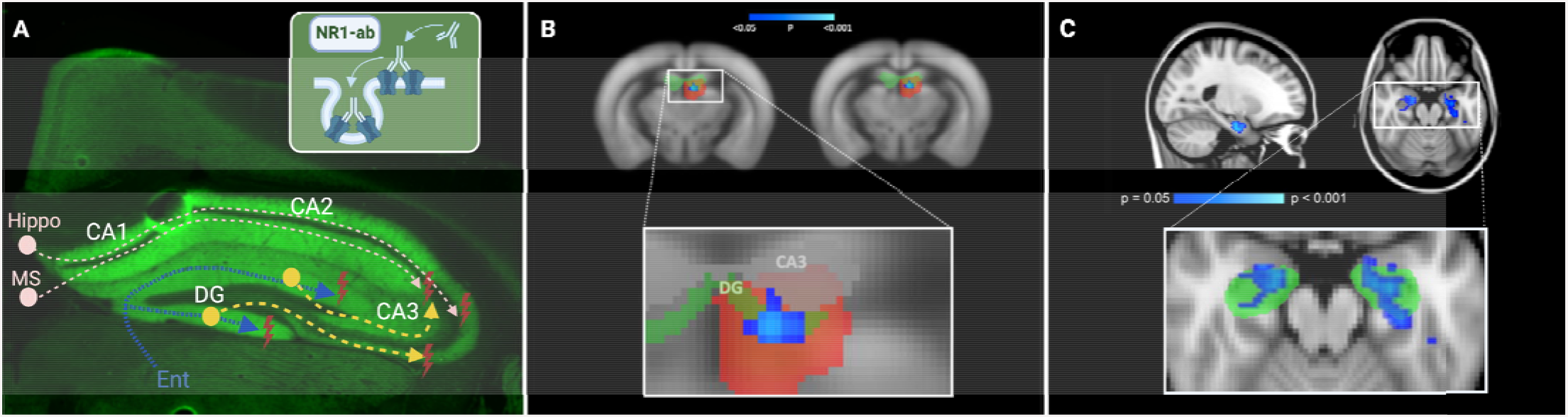
Comparative pathology and MR correlates of impaired hippocampal functional connectivity in human anti-NMDAR encephalitis and murine NMDAR antibody disease models. **A**: Pathophysiological mechanisms in murine NMDAR encephalitis disease model, drawn in the immunofluorescence staining of a coronal brain section from a mouse of cohort 1. NR1-antibodies lead to degradation of hippocampal NMDA receptors in dentate gyrus and CA1/CA3/CA3 (insert). Projection fibers from entorhinal cortex (blue) are entering *stratum moleculare* of dentate gyrus via perforant path. Pyramidal neurons in CA3 receive intrahippocampal projection fibers from dentate gyrus (yellow) and from external fibers from medial septum and contralateral hippocampus (pink). Dysfunction of hippocampal neurons in DG and CA3 lead to impairment in the connectivity of hippocampal circuits. **B:** Reduced functional connectivity (significantly different area displayed based on p-values in blue-lightblue) within hippocampus (hippocampal functional component area displayed as z-map in red-yellow; anatomical area of dentate gyrus displayed in green) of NR1 mice compared to controls (cohort 1), demonstrated using the same coronal plane as in A. **C:** Reduced functional connectivity (blue) between both the left and the right hippocampus (green area) and the anterior default-mode network in NMDARE patients compared to healthy controls (original picture published in ^4^). Ent = entorhinal cortex; NR1-ab = NR1 IgG antibodies; NMDARE = anti-NMDAR encephalitis; MS = medial septum; Hippo = contralateral hippocampus; DG = dentate gyrus; Sub = subiculum

## Discussion

We observed selectively impaired functional connectivity of the hippocampus in an adult mouse model of NMDAR encephalitis. In addition, selective impairment of hippocampal functional connectivity was found in mice with *in utero* exposure to monoclonal NR1 antibodies at two different time points. These findings mirror observations of functional connectivity alterations in human patients with NMDARE and are in line with the predominant expression of NMDAR in the hippocampus (Figure 5 B and C). As such, our study identifies the first cross-species imaging biomarker in NMDAR encephalitis and provides a promising foundation for translational studies on the pathophysiology and on preclinical treatment evaluation of the disease.

### Comparison of mouse data with human data

In a whole brain functional connectivity analysis in a mouse model of NMDAR encephalitis, we observed a selective disruption of hippocampal connectivity that is remarkably consistent with findings in human patients: In a sample of 24 NMDAR encephalitis patients, selectively reduced functional connectivity of the hippocampus with the anterior default mode network (DMN) was found and the level of hippocampal connectivity disruption correlated with the severity of memory impairment ^4^. Subsequent large-scale network and whole-brain pair-wise connectivity analyses in 43 NMDAR encephalitis patients corroborated this finding of impaired hippocampal connectivity alongside decoupling of medial temporal network and the DMN and an overall impairment of frontotemporal connections that correlated with memory impairment and schizophrenia-like symptoms ^5^. Our current results of selectively impaired functional connectivity of the hippocampus in mice after passive transfer of NMDAR antibodies indicate (i) the central role of hippocampal dysfunction in the pathophysiology of NMDAR encephalitis, (ii) the validity of the employed NMDAR encephalitis mouse model mirroring the findings in human patients, (iii) the relevance of materno-fetal NMDAR antibody transmission and (iv) the translational applicability of resting state functional MRI analyses – as will be discussed in the following.

#### (i) Central role of hippocampal dysfunction

The predominant vulnerability of the hippocampus in NMDAR encephalitis is in line with the fact that hippocampus contains the highest density of NMDARs in the human brain ^6^. Particularly, the NR1 subunit of the NMDAR is widely expressed in all parts of the human hippocampus, including the dentate gyrus, CA1-CA3, subiculum, presubiculum and entorhinal cortex ^30^. Similar distribution patterns are seen in rodent brains with hippocampal levels of NR1 protein widely exceeding the levels of cortex, olfactory bulb, midbrain and cerebellum ^31^. Volumetric MRI analyses in human patients with NMDAR encephalitis lent further support to this concept of predominant hippocampal damage by showing localized atrophy of hippocampal subfields that correlated with individual deficits in verbal memory performance ^32^.

However, the clinical syndrome of NMDAR encephalitis clearly extends beyond hippocampal dysfunction and suggests additional cortical and subcortical involvement ^33^. This is supported by recent studies showing reduced functional connectivity in distributed large-scale networks, including sensorimotor, frontoparietal, and visual networks ^5^ and is in line with global metabolic changes observed using positron emission tomography (PET) ^34,35^. In addition, extensive white matter damage related to disease severity and cognitive impairment was observed in NMDAR encephalitis patients using diffusion-weighted imaging ^4,36^. Together, these studies indicate brain-wide dysfunction of NMDA receptors, with predominant alteration of hippocampal function. Indeed, NMDAR antibodies were shown to bind throughout the rodent brain - with strong preferential binding in the hippocampus, leading to decreased NMDAR cluster density on neurons ^37^. In addition, it was recently found that NMDAR antibodies also alter the function of NMDAR in oligodendrocytes, thus providing a direct link between antibody-mediated NMDAR dysfunction and white matter damage observed using MRI ^38^. It is therefore conceivable that future studies with further refined animal models and imaging protocols will be able to detect additional functional connectivity changes as predicted by human MRI studies. Furthermore, diffusion tensor imaging studies – ideally in combination with histopathological analyses – are warranted to investigate white matter integrity in NMDAR encephalitis mouse models to advance the understanding of oligodendrocyte pathophysiology in this disease.

#### (ii) NMDAR encephalitis mouse models

In our NMDAR encephalitis model, mice were infused with a human monoclonal IgG1 antibody into the lateral ventricles which strongly bound to NMDA receptors in the brain. While the animals do not develop an obvious clinical phenotype, such as movement disorders or overt seizures, the seizure threshold is reduced *in vivo* ^11^, recently shown to result from reduced synaptic excitatory neurotransmission related to alterations in the dynamical behavior of brain microcircuits ^39^. Thus, passive immunization of rodents with human monoclonal NR1 antibodies resulted in similar electrophysiology changes as in humans. Likewise, behavioral alterations in human patients and mice showed remarkable similarities, both in passive immunization using enriched human IgG from NMDAR encephalitis patients ^8^ and active immunization with NMDAR holoreceptors ^7^ or receptor peptides ^9,10^. The here observed consistent finding of impaired hippocampal connectivity in mice and in humans further suggests that NR1 autoantibodies mediate disease mechanisms in a similar way across species. Further research is needed to determine whether active immunization ^7^ may better mimic the human disease compared to passive antibody transfer performed here, which might limit transferability from our findings to human disease pathology. However, as electrophysiological and functional brain changes occur down-stream of the antibody binding, passive immunization likely provides a meaningful model to study fMRI changes across species.

#### (iii) Mouse model of NR1 antibody exposure *in utero*

Functional connectivity analyses were additionally carried out in a recently established murine model with *in utero* exposure to monoclonal NR1 antibodies ^14^. Very similar functional connectivity reductions of the hippocampus were observed in offspring at the age of 8 weeks and 10 months. Transferred human gestational NMDAR autoantibodies have recently been shown to accumulate in the fetus leading to a considerable reduction of synaptic NMDAR density in the early neonate mouse brain. Adult NR1 offspring animals displayed an abnormal neuropsychiatric phenotype, including hyperactivity, lower anxiety and impaired sensorimotor gating that was complemented by structural MRI atrophy of cerebral cortex and brainstem ^14^. It was therefore suggested that diaplacentally transferred NR1 antibodies could potentially contribute to a broader spectrum of behavioral abnormalities found in autism spectrum disorders, bipolar disorders or schizophrenia, thus diverging from typical characteristics of adult NR1-antibody mediated disease ^14^. Our findings provide evidence for a long-term impairment of hippocampal development and function caused by fetal NR1-antibody exposure, reflected by a predominant reduction in hippocampal functional connectivity in adult offspring mice. These findings complement structural brain changes with whole brain and regional atrophy in the same offspring NR1 mouse model that were linked to a possible NR1-antibody mediated neurodegenerative mechanism associated with a clinical phenotype. While functional connectivity alterations in other parts of the brain were not detected, future studies combining functional and structural MR data with neuropsychiatric features will help to further elucidate the pathophysiological mechanisms in this gestational mouse model of maternal NMDAR antibodies. In the future, similar studies should be expanded to other autoantibodies, e.g. Caspr2, for which a neurodevelopmental phenotype after materno-fetal antibody transfer was recently suggested ^40^.

#### (iv) Translational applicability of resting-state fMRI

Preclinical investigations using rs-fMRI in murine disease models offer the possibility to non-invasively investigate the determinants of altered functional network signatures observed in human studies, including associations with behavioral changes, structural MRI damage and histopathological correlates ^16^. Despite limiting factors inherent to mouse rs-fMRI at 7 Tesla, including effects of anesthesia on brain activity and vasculature ^41^, and challenges in maintaining constant animal physiology throughout the entire MR investigation ^42^, mouse rs-fMRI has become a reliable and valuable tool to unravel pathophysiological mechanisms in neuropsychiatric diseases – in part due to improvement of anesthesia protocols and continuous vital parameter monitoring including temperature and breathing rate measurement as performed in our study. Previous studies have shown highly reproducible large-scale networks in multi-centre comparisons ^16^ and have provided promising accounts of translational functional network analyses in neuropsychiatric and neuroinflammatory diseases such as Alzheimer’s disease ^43^, depression ^44^, schizophrenia ^45^ and demyelinating disorders ^46^ (for further extensive reviews see Chuang and Nasrallah 2017 and Pan et al. 2015 ^41,42^). These observations add to the growing translational evidence of functional network alterations in neuropsychiatric diseases and provide a promising avenue for future comparative studies in NMDAR encephalitis patients and animal models of the disease. Aims of this approach include the identification of biomarkers for the assessment of preclinical treatment responses as well as comparative analyses of MRI alterations with histopathologic findings.

In line with previous studies that investigated rs-fMRI in mouse models in other neuropsychiatric disorders such as Huntington’s disease ^47^ and Fragile-X syndrome ^48^, our findings suggest that functional connectivity may eventually allow for non-invasive monitoring of disease activity and effects of pharmacological interventions in NR1 autoimmune encephalitis mouse models. Hence, rs-fMRI functional connectivity, as a surrogate marker for brain network integrity, could complement behavioral assays – which suffer from limited translational validity – in future drug development in rodents ^48^.

## Conclusion

Our study demonstrates the feasibility and validity of fMRI-based functional connectivity analyses in NMDAR antibody-mediated murine disease models and reveals a selectively impaired functional connectivity of the hippocampus. This finding mirrors observations in human patients and represents the first translational imaging biomarker of NMDAR encephalitis. Our investigations in mice with *in utero* exposure to NR1 antibodies underline the possibility to experimentally explore antibody-mediated mechanisms in constellations that are only limitedly or not at all accessible to human studies. Furthermore, our study provides an important foundation for future comparative structural and functional MRI analyses in monoclonal NR1 antibody-based animal models and human patients, including interventional studies. These investigations will combine structural and functional MRI analyses, including detailed regional and large-scale network analyses, with histopathologic studies to further unravel the pathophysiology of NMDAR encephalitis, e.g. the mechanisms involved in hippocampal functional connectivity alterations and in structural (e.g. white matter) brain damage.

## Acknowledgments

We thank Claudia Sommer for her excellent technical support. JKu and JKr are participants in the BIH-Charité (Junior) Clinician Scientist Program funded by the Charité – Universitätsmedizin Berlin and the Berlin Institute of Health.

## References

1. Graus, F. et al. A clinical approach to diagnosis of autoimmune encephalitis. Lancet Neurol. 15, 391–404 (2016).

2. Dalmau, J., Lancaster, E., Martinez-Hernandez, E., Rosenfeld, M. R. & Balice-Gordon, R. Clinical experience and laboratory investigations in patients with anti-NMDAR encephalitis. Lancet Neurol. 10, 63–74 (2011).

3. Heine, J. et al. Imaging of autoimmune encephalitis--Relevance for clinical practice and hippocampal function. Neuroscience 309, 68–83 (2015).

4. Finke, C. et al. Functional and structural brain changes in anti-N-methyl-D-aspartate receptor encephalitis. Ann. Neurol. 74, 284–296 (2013).

5. Peer, M. et al. Functional connectivity of large-scale brain networks in patients with anti-NMDA receptor encephalitis: an observational study. Lancet Psychiatry 4, 768–774 (2017).

6. Monaghan, D. T. & Cotman, C. W. Distribution of N-methyl-D-aspartate-sensitive L-[3H]glutamate-binding sites in rat brain. J. Neurosci. Off. J. Soc. Neurosci. 5, 2909–2919 (1985).

7. Jones, B. E. et al. Autoimmune receptor encephalitis in mice induced by active immunization with conformationally stabilized holoreceptors. Sci. Transl. Med. 11, (2019).

8. Planagumà, J. et al. Human N-methyl D-aspartate receptor antibodies alter memory and behaviour in mice. Brain J. Neurol. 138, 94–109 (2015).

9. Wagnon, I. et al. Autoimmune encephalitis mediated by B-cell response against N-methyl-d-aspartate receptor. Brain 143, 2957–2972 (2020).

10. Wilke, J. B. H. et al. Autoantibodies against NMDA receptor 1 modify rather than cause encephalitis. Mol. Psychiatry (2021) doi:10.1038/s41380-021-01238-3.

11. Wright, S. et al. Epileptogenic effects of NMDAR antibodies in a passive transfer mouse model. Brain 138, 3159–3167 (2015).

12. Kreye, J. et al. Human cerebrospinal fluid monoclonal N-methyl-D-aspartate receptor autoantibodies are sufficient for encephalitis pathogenesis. Brain J. Neurol. 139, 2641–2652 (2016).

13. Wenke, N. K. et al. N-methyl-D-aspartate receptor dysfunction by unmutated human antibodies against the NR1 subunit. Ann. Neurol. 85, 771–776 (2019).

14. Jurek, B. et al. Human gestational N-methyl-d-aspartate receptor autoantibodies impair neonatal murine brain function. Ann. Neurol. 86, 656–670 (2019).

15. García-Serra, A. et al. Placental transfer of NMDAR antibodies causes reversible alterations in mice. Neurol. Neuroimmunol. Neuroinflammation 8, e915 (2021).

16. Grandjean, J. et al. Common functional networks in the mouse brain revealed by multi-centre resting-state fMRI analysis. bioRxiv 541060 (2019) doi:10.1101/541060.

17. Mechling, A. E. et al. Fine-grained mapping of mouse brain functional connectivity with resting-state fMRI. NeuroImage 96, 203–215 (2014).

18. Sert, N. P. du et al. The ARRIVE guidelines 2.0: Updated guidelines for reporting animal research. PLOS Biol. 18, e3000410 (2020).

19. Ly, L.-T. et al. Affinities of human NMDA receptor autoantibodies: implications for disease mechanisms and clinical diagnostics. J. Neurol. 265, 2625–2632 (2018).

20. Kreye, J. et al. Encephalitis patient-derived monoclonal GABAA receptor antibodies cause epileptic seizures. J. Exp. Med. 218, e20210012 (2021).

21. Petit-Pedrol, M. et al. LGI1 antibodies alter Kv1.1 and AMPA receptors changing synaptic excitability, plasticity and memory. Brain J. Neurol. 141, 3144–3159 (2018).

22. Planagumà, J. et al. Ephrin-B2 prevents N-methyl-D-aspartate receptor antibody effects on memory and neuroplasticity. Ann. Neurol. 80, 388–400 (2016).

23. Wardemann, H. et al. Predominant autoantibody production by early human B cell precursors. Science 301, 1374–1377 (2003).

24. Zerbi, V., Grandjean, J., Rudin, M. & Wenderoth, N. Mapping the mouse brain with rs-fMRI: An optimized pipeline for functional network identification. NeuroImage 123, 11–21 (2015).

25. Jenkinson, M., Bannister, P., Brady, M. & Smith, S. Improved optimization for the robust and accurate linear registration and motion correction of brain images. NeuroImage 17, 825–841 (2002).

26. Beckmann, C. F. & Smith, S. M. Probabilistic independent component analysis for functional magnetic resonance imaging. IEEE Trans. Med. Imaging 23, 137–152 (2004).

27. Griffanti, L. et al. Hand classification of fMRI ICA noise components. Neuroimage 154, 188–205 (2017).

28. Filippini, N. et al. Distinct patterns of brain activity in young carriers of the APOE-epsilon4 allele. Proc. Natl. Acad. Sci. U. S. A. 106, 7209–7214 (2009).

29. Ceccom, J., Halley, H., Daumas, S. & Lassalle, J. M. A specific role for hippocampal mossy fiber’s zinc in rapid storage of emotional memories. Learn. Mem. 21, 287–297 (2014).

30. Scherzer, C. R. et al. Expression of N-methyl-D-aspartate receptor subunit mRNAs in the human brain: hippocampus and cortex. J. Comp. Neurol. 390, 75–90 (1998).

31. Luo, J., Bosy, T. Z., Wang, Y., Yasuda, R. P. & Wolfe, B. B. Ontogeny of NMDA R1 subunit protein expression in five regions of rat brain. Brain Res. Dev. Brain Res. 92, 10–17 (1996).

32. Finke, C. et al. Structural Hippocampal Damage Following Anti-N-Methyl-D-Aspartate Receptor Encephalitis. Biol. Psychiatry 79, 727–734 (2016).

33. Dalmau, J. et al. An update on anti-NMDA receptor encephalitis for neurologists and psychiatrists: mechanisms and models. Lancet Neurol. 18, 1045–1057 (2019).

34. Leypoldt, F. et al. Fluorodeoxyglucose positron emission tomography in anti-N-methyl-D-aspartate receptor encephalitis: distinct pattern of disease. J. Neurol. Neurosurg. Psychiatry 83, 681–686 (2012).

35. Probasco, J. C. et al. Decreased occipital lobe metabolism by FDG-PET/CT: An anti-NMDA receptor encephalitis biomarker. Neurol. Neuroimmunol. Neuroinflammation 5, e413 (2018).

36. Phillips, O. R. et al. Superficial white matter damage in anti-NMDA receptor encephalitis. J. Neurol. Neurosurg. Psychiatry 89, 518–525 (2018).

37. Moscato, E. H. et al. Acute mechanisms underlying antibody effects in anti-N-methyl-D-aspartate receptor encephalitis. Ann. Neurol. 76, 108–119 (2014).

38. Matute, C. et al. N-Methyl-D-Aspartate Receptor Antibodies in Autoimmune Encephalopathy Alter Oligodendrocyte Function. Ann. Neurol. 87, 670–676 (2020).

39. Wright, S. K. et al. Multimodal electrophysiological analyses reveal that reduced synaptic excitatory neurotransmission underlies seizures in a model of NMDAR antibody-mediated encephalitis. Commun. Biol. 4, 1106 (2021).

40. Coutinho, E. et al. Persistent microglial activation and synaptic loss with behavioral abnormalities in mouse offspring exposed to CASPR2-antibodies in utero. Acta Neuropathol. (Berl.) 134, 567–583 (2017).

41. Chuang, K.-H. & Nasrallah, F. A. Functional networks and network perturbations in rodents. NeuroImage 163, 419–436 (2017).

42. Pan, W.-J., Billings, J. C. W., Grooms, J. K., Shakil, S. & Keilholz, S. D. Considerations for resting state functional MRI and functional connectivity studies in rodents. Front. Neurosci. 9, 269 (2015).

43. Grandjean, J. et al. Early alterations in functional connectivity and white matter structure in a transgenic mouse model of cerebral amyloidosis. J. Neurosci. Off. J. Soc. Neurosci. 34, 13780–13789 (2014).

44. Grandjean, J. et al. Chronic psychosocial stress in mice leads to changes in brain functional connectivity and metabolite levels comparable to human depression. NeuroImage 142, 544–552 (2016).

45. Gass, N. et al. An acetylcholine alpha7 positive allosteric modulator rescues a schizophrenia-associated brain endophenotype in the 15q13.3 microdeletion, encompassing CHRNA7. Eur. Neuropsychopharmacol. J. Eur. Coll. Neuropsychopharmacol. 26, 1150–1160 (2016).

46. Hübner, N. S. et al. The connectomics of brain demyelination: Functional and structural patterns in the cuprizone mouse model. NeuroImage 146, 1–18 (2017).

47. Li, Q. et al. Resting-state functional MRI reveals altered brain connectivity and its correlation with motor dysfunction in a mouse model of Huntington’s disease. Sci. Rep. 7, 16742 (2017).

48. Zerbi, V. et al. Inhibiting mGluR5 activity by AFQ056/Mavoglurant rescues circuit-specific functional connectivity in Fmr1 knockout mice. NeuroImage 191, 392–402 (2019).

49. Mechling, A. E. et al. Deletion of the mu opioid receptor gene in mice reshapes the reward-aversion connectome. Proc. Natl. Acad. Sci. U. S. A. 113, 11603–11608 (2016).

